# Structural basis of recognition and destabilization of histone H2B ubiquitinated nucleosome by DOT1L histone H3 Lys79 methyltransferase

**DOI:** 10.1101/508663

**Authors:** Seongmin Jang, Chanshin Kang, Han-Sol Yang, Taeyang Jung, Hans Hebert, Ka Young Chung, Seung Joong Kim, Sungchul Hohng, Ji-Joon Song

## Abstract

DOT1L is a histone H3 Lys79 methyltransferase whose activity is stimulated by histone H2B Lys120 ubiquitination, suggesting cross-talk between histone H3 methylation and H2B-ubiquitination. Here, we present cryo-EM structures of DOT1L complex with unmodified and H2B-ubiquitinated nucleosomes, showing that DOT1L recognizes H2B-ubiquitin and the H2A/H2B acidic patch through a C-terminal hydrophobic helix and an arginine anchor in DOT1L respectively. Furthermore, the structures combined with single-molecule FRET experiment show that H2B-ubiquitination enhances a non-catalytic function of DOT1L destabilizing nucleosome. These results establish the molecular basis of the cross-talk between H2B ubiquitination and H3 Lys79 methylation as well as nucleosome destabilization by DOT1L.

## Introduction

The fundamental unit of chromatin is the nucleosome, a structure composed of a histone octamer and 146 bp DNA (Luger et al. 1997). The structure of the nucleosome is altered by ATP-dependent chromatin remodelers and histone chaperones. A recent cryo-EM work showed that the nucleosome has structural flexibility as DNA breathing leads to conformational changes in the histones (Bilokapic et al. 2018). The histones in the nucleosome are covalently modified with many moieties including methyl, acetyl, phosphor, and ubiquitin (Gurard-Levin et al. 2014; Lawrence et al. 2016; Clapier et al. 2017). These histone covalent modifications are key mechanisms that modulate epigenetic gene expression. Among these, histone methylation is very complex as the locations and levels of histone methylations lead to different consequences in gene expression. For example, histone H3 Lys4 methylation by MLL methyltransferase is a hallmark for gene activation, while histone H3 Lys27 methylation by PRC2 methyltransferase is a marker for gene repression (Lawrence et al. 2016). While most histone methylations occur in the N-terminal flexible tails in histones, histone H3 Lys 79 is located on the body of the nucleosome. Histone H3 Lys79 methylation is highly correlated with actively transcribing genes and occurs by the methyltransferase Yeast Dot1 and human DOT1L, enzymes that have non-SET methyltransferase domains (Min et al. 2003; Sawada et al. 2004; Nguyen and Zhang 2011). Histone methylation has another complexity, as some methylations are regulated by other histone modifications suggesting a cross-talk between modifications (Lee et al. 2010). Mono-ubiquitination of histone H2B Lys120 was shown to be required for H3 Lys79 methylation *in-vitro* and *in-vivo* (Briggs et al, McGinty et al, Ng et al). These findings suggest that there is crosstalk between H2B ubiquitination and histone H3 Lys79 methylation. However, it is not clear how DOT1L recognizes H2B-ubiquitinated nucleosomes, thus leading to stimulation of DOT1L activity.

In addition to their catalytic activities, histone methyltransferases have been shown to have non-catalytic activities (Kim et al. 2015). A recent study showed that yeast Dot1 has nucleosome chaperone activity and enhances the ATP-dependent chromatin remodeling by CHD1, which is independent of Dot1 methyltransferase activity (Lee et al. 2018). This work suggested that Dot1/DOT1L might destabilize nucleosome structure and help the remodeler to alter nucleosome structure. However, the molecular details of DOT1L mediated nucleosome destabilization have not been investigated.

Here, we present cryo-EM structures of DOT1L bound to unmodified or H2B-ubiquitinated nucleosomes. The structures revealed that the relative location of DOT1L to nucleosome is reoriented in ubiquitinated nucleosome (Nuc_H2B-Ub) compared with unmodified nucleosome (Nuc), and that DOT1L recognizes H2B-Ub via two main modes; a hydrophobic helix located at the C-terminal of DOT1L catalytic domain recognizes H2B-Ub. In addition, the cryo-EM structures reveal that DNA is detached from the histones and the first helix of H3 and the C-terminus of H2A are concomitantly disordered, indicating that DOT1L binding destabilizes the nucleosome structure. Furthermore, single-molecule FRET analysis shows that H2B ubiquitination greatly increases DOT1L-mediated nucleosome destabilization suggesting that H2B ubiquitination affects not only histone H3 methylation but also nucleosome destabilization by DOT1L. Overall, this work provides the first molecular insight into a cross-talk between H2B-Ub and H3 Lys79 methylation as well as a structural view of DOT1L mediated nucleosome destabilization.

## Results and Discussion

### Cryo-EM structures of DOT1L-Nucleosome with or without ubiquitin

DOT1L methylates histone H3 Lys79 located on the body of nucleosome and its activity is stimulated by H2B-ubiquitin. To understand how DOT1L recognizes ubiquitinated nucleosomes, we determined cryo-EM structures of DOT1L catalytic domain (1-416 a.a.) bound to unmodified or H2B-ubiquitinated nucleosomes. We prepared ubiquitinated histone H2B (H2B-Ub) by chemical crosslinking (Morgan et al. 2016), and assembled H2B-Ub nucleosome by mixing ubiquitinated histone octamers with 601 DNA (Lowary and Widom 1998) (Supplemental Figs. 1A and 1B). We then purified DOT1L_nucleosomes (DOT1L_Nuc) and DOT1L_H2B-ubiquitinated nucleosomes (DOT1L_Nuc_H2B-Ub) by GraFix (Stark 2010). Micrographs from plunge-frozen DOT1L-nucleosome complexes were collected using Titan Krios 300KeV with a Gatan K2 Summit direct detect (Supplemental Table 1). The micrographs were processed and refined with Relion2 (Kimanius et al. 2016). The FSC of the cryo-EM maps indicates 6.8 Å or 7.3 Å resolutions for DOT1L_Nuc-H2B_Ub and for DOT1L_Nuc_H2B-Ub respectively (Supplemental Figs. 2 and 3). Despite of the relatively low resolutions, the secondary structures of histones and the grooves of DNA were well resolved (Figs. 1A, 1B and Supplemental Fig. 4). The cryo-EM maps clearly indicated that DOT1L is bound to only one side of the nucleosome. The cryo-EM map of DOT1L is relatively weak, suggesting that the binding of DOT1L to the nucleosome is flexible and that DOT1L might have multiple binding modes to the nucleosome. Comparison between the cryo-EM maps of DOT1L_Nuc and DOT1L_Nuc_H2B-Ub clearly located the H2B ubiquitin (Figs. 1A, 1B and 1E).

**Figure 1.**
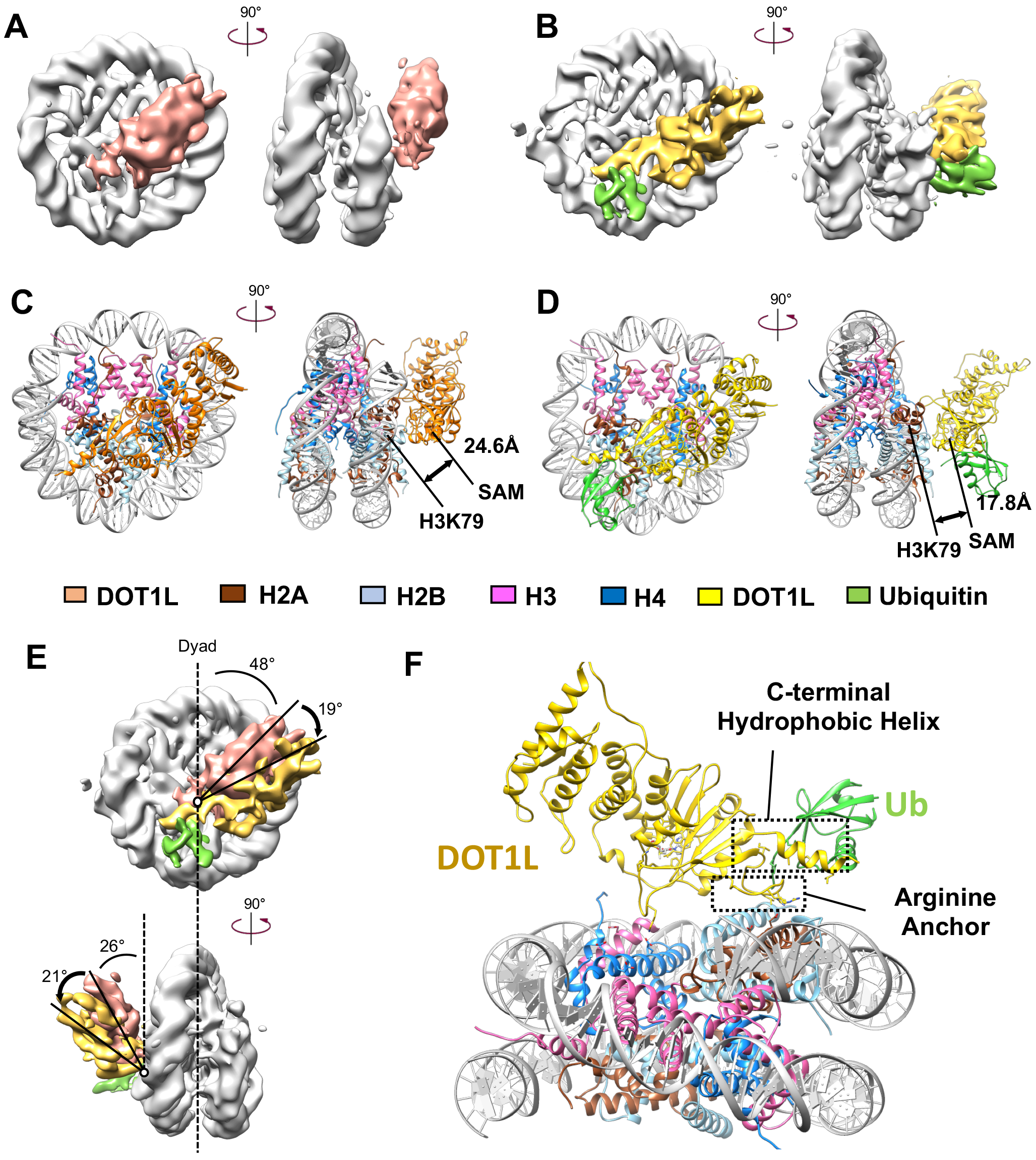
Cryo-EM structures of DOT1L_nucleosomes. (A) Cryo-EM map of DOT1L bound to unmodified nucleosome (DOT1L_Nuc). Nucleosome is shown in gray, and DOT1L in pale red. The map of DOT1L was drawn in lower contour than nucleosome due to the week map of DOT1L. (B) Cryo-EM structure of DOT1L bound to H2B ubiquitinated nucleosome (Nuc_H2B-Ub). Nucleosome is shown in gray, DOT1L in yellow, and ubiquitin in green. (C) Atomic model of DOT1L_Nuc is shown in two orientations. (D) Atomic model of DOT1L_Nuc_H2B-Ub is shown in two orientations. (E) Superimpositions of cryo-EM maps of DOT1L_Nuc and DOT1L_Nuc_H2B-Ub. The axes for measuring the relative rotation angles from the two-fold axis of nucleosome are in dotted lines. (F) An enlarged view of the DOT1L_Nuc_H2B-Ub. The location of the C-terminal hydrophobic helix and the arginine anchor are indicated with the dotted boxes.

We generated models of DOT1L_Nuc and DOT1L_Nuc_H2B-Ub by fitting the crystal structures of DOT1L (PDB ID: 1NW3), the nucleosome (PDB ID: 1KX3), and ubiquitin (PDB ID: 1UBI) and the models were further refined with molecular dynamics flexible fitting (MDFF) (Ramage et al. 1994; Davey et al. 2002; Min et al. 2003; Trabuco et al. 2008) (Figs. 1C, 1D and Supplemental Fig. 4). In the DOT1L_Nuc complex, DOT1L bound to the nucleosome rotated in the clock-wise by 48 degrees from the two fold axis of the nucleosome (Fig. 1E). The C-terminal half of DOT1L containing the catalytic pocket is located near the target histone H3 Lys79. However, the N-terminal half of DOT1L interacts with neither DNA nor histones. In this configuration, the region near the catalytic pocket is attached to the nucleosome and the N-terminal half is detached from the nucleosome with 26 degrees from the symmetric axis of the nucleosome from the side-view. In the DOT1L_Nuc_H2B-Ub complex, the overall binding mode of DOT1L is overall similar to the DOT1L_Nuc complex (Fig. 1B). Superimposition between cryo-EM maps of DOT1L_Nuc and DOT1L_Nuc_H2B-Ub revealed that the DOT1L with H2B-Ub nucleosome is further rotated about 19 degrees and the N-terminal half is also further detached from nucleosome in with 21 degrees compared with DOT1L_Nuc (Figs. 1E and 1F).

The distances between S-Adenosyl Methionines (SAMs) and the target histone H3 Lys79 are 24.6 Å for DOT1L_Nuc and 17.8 Å for DOT1L_Nuc_H2B-Ub (Figs. 1C and 1D). The closer distance of DOT1L to Nuc_H2B-Ub nucleosome than DOT1L to Nuc suggests that the H2B-Ub might reorient DOT1L toward the target histone H3 Lys79. However, the distances between SAM and H3 Lys79 in both structures are too far to methylate the target, suggesting that the current structures might be catalytically inactive states where the substrate is not fully engaged. These results show that the C-terminal part of DOT1L catalytic domain interacts with nucleosome while the N-terminal part is detached from the nucleosome and that DOT1L might have multiple binding orientations, suggesting that H2B-ubiquitination might reorient DOT1L into a more active orientation.

### The recognition of H2B-ubiquitin by DOT1L via a hydrophobic C-terminal helix

Histone H3 Lys79 methylation by DOT1L is stimulated by H2B-ubiquitination, indicating the direct interaction between DOT1L and the ubiquitin. Unfortunately, in our DOT1L_Nuc_H2B-Ub structure, the exact rotational position of the ubiquitin could not be resolved due to the poor map of the ubiquitin. However, the cryo-EM structures clearly show the close proximity of H2B-ubiquitin and the C-terminal helix of DOT1L catalytic domain (Fig. 2A and Supplemental Fig. 4B). A previous mutational analysis on the surface of ubiquitin revealed that the hydrophobic surface near L71 and L73 located at the C-terminal of ubiquitin is critical for DOT1L activity (Holt et al. 2015).

**Figure 2.**
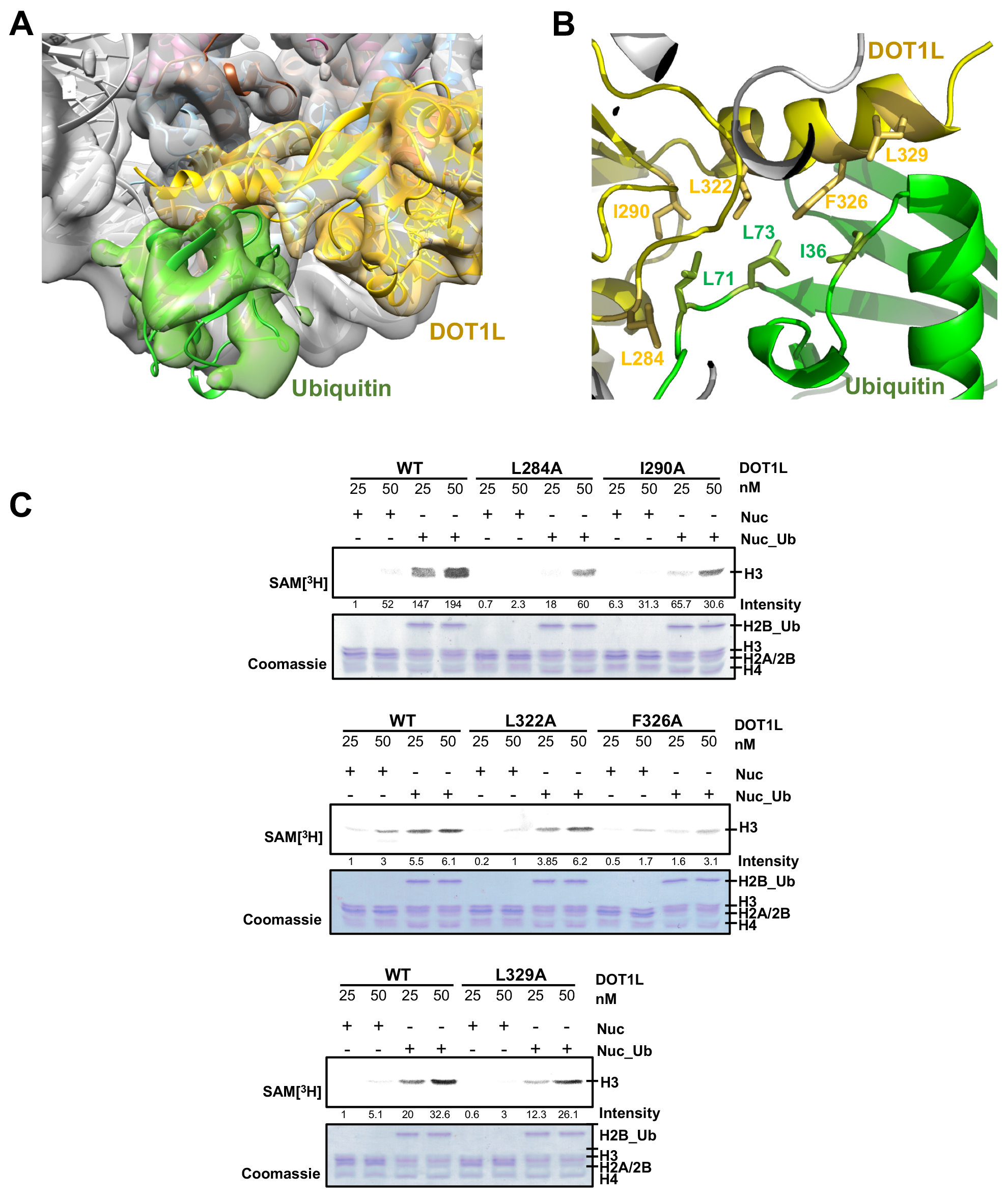
DOT1L recognizes the H2B Lys120 ubiquitin via the C-terminal hydrophobic helix. (A) Cryo-EM map with the model near the C-terminal hydrophobic helix. (B) Detailed interaction between the C-terminal hydrophobic helix and H2B-ubiquitin. (C) Histone methyltransferase assay with wild-type and the mutants where the hydrophobic residues (L284, I290, L322, F325, L329) were mutated to alanines. Autoradiographs for methylated histone H3 Lys79 with ^3^H-SAM (top panel). The half of each reaction was run in a separate gel and stained with Coomassie blue for a loading control (bottom panel). Intensities of the methylated histones were measured and written in numbers under the gel.

The C-terminal helix (321-332 a.a.) of the DOT1L catalytic domain exhibits an amphipathic character where one side of the helix is hydrophobic and the other side is hydrophilic. Several hydrophobic residues (L322A, F326A and L329A) in the helix together with hydrophobic residues (L284 and L290) in the loop near the helix forms a hydrophobic patch (Figs. 2A and 2B). Interestingly, in our model, the hydrophobic patch in DOT1L is facing the previously identified hydrophobic surface including L71 and L73 in the ubiquitin in H2B, suggesting that DOT1L might recognize H2B-ubiquitin via the hydrophobic C-terminal helix. Specifically, L322, F326 and L329 seem to interact with the previously identified hydrophobic residues (L71 and L73) together with I36 in H2B-ubiquitin, and L284 is located near the C-terminus of the ubiquitin (Fig. 2B). To determine whether these hydrophobic residues in DOT1L are critical for DOT1L activity, we mutated these hydrophobic residues to alanines (L284A, I290A, L322A, F326A and L329A) and examined the DOT1L HMTase activity (Fig. 2C and Supplemental Fig. 1C). Except L322A, all alanine mutants significantly decreased DOT1 activity. These results reveal that DOT1L recognizes H2B-Ub via the C-terminal hydrophobic helix of DOT1L and these interactions are important for DOT1L HMTase activity.

### DOT1L recognizes the H2A/H2B acidic patch in nucleosome via an arginine anchor

Several nucleosome binding proteins use arginine anchors to interact with the H2A/H2B acidic patch (Makde et al. 2010; Armache et al. 2011; Kato et al. 2013; Morgan et al. 2016). Interestingly, two conserved arginines (R278 and R282) exist in the loop between amino acid 268 and 286 in DOT1L. The cryo-EM structure shows that these arginines recognize the H2A/H2B acidic patch, suggesting that DOT1L might also utilize the arginine anchor to recognize the acidic patch in the nucleosome (Figs. 3A and 3B). To examine the role of the arginine anchor for DOT1L activity, we mutagenized the arginines to alanines (R278A and R282A) and measure DOT1L HMTase activities (Fig. 3C and Supplemental Fig. 1C). The structure shows that R282 seems to directly interacts with E61 and E63 from H2A, and E110 from H2B and R278 is located closed to these glutamates (Fig. 3B). The arginine mutant DOT1L_R282A_ but not DOT1L_R278A_ dramatically decreased DOT1L activity, implying that DOT1L also utilizes the arginine anchor to interact with nucleosome as other nucleosome binding proteins.

**Figure 3.**
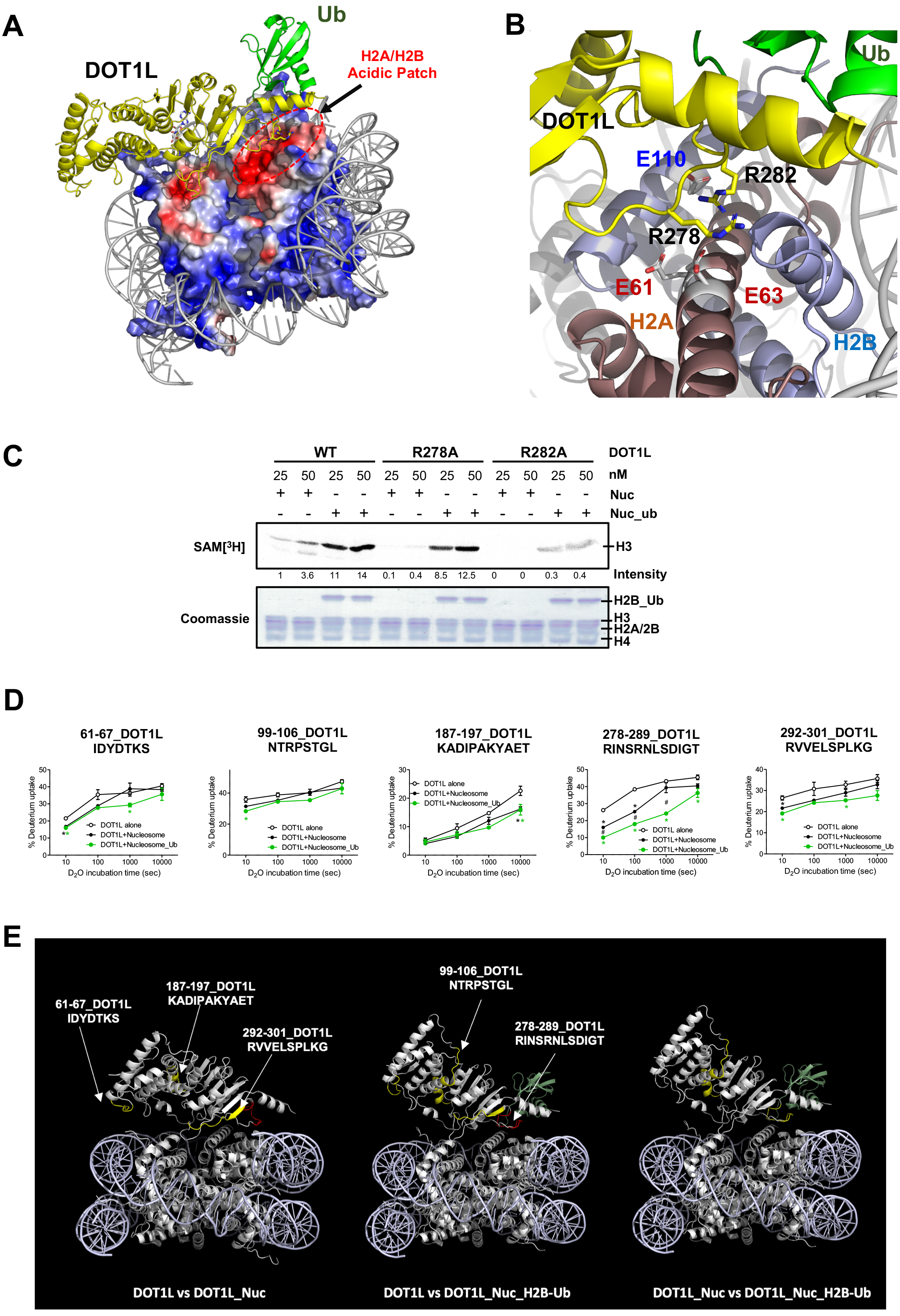
DOT1L recognizes the H2A/H2B acidic patch with the arginine anchor. (A) The arginine anchor is located to near the H2A/H2B acidic patch. (B) Detailed view for the interaction between the arginines (R278 and R282) in the arginine anchor and the glutamates (E61h2a, E63h2a and E110h2b) in the H2A/H2B acidic patch. (C) DOT1L HMTase assay with wild-type and the arginine anchor mutants (R278A and R282A). Autoradiographs for methylated histone H3 Lys79 with ^3^H-SAM (top panel). The half of each reaction was run in a separate gel and stained with Coomassie blue for a loading control (bottom). Intensities of the methylated histones were measured and written in numbers under the gel. (D) The deuterium uptake plots of selected DOT1L peptides of DOT1L showing the difference in deuterium uptake rate between states. The significant difference (P < 0.05) were indicated as * between DOT1L alone and DOT1L+ nucleosome, * (green) between DOT1L alone and DOT1L+H2B-ubiquitinated nucleosome, and # between DOT1L + nucleosome and DOT1L+H2B-ubiquitinated nucleosome. (E) The peptides analyzed in HDX-MS were shown in the structures. The peptides colored in red indicates higher difference in the deuterium uptakes than the peptides in yellow. The arginine anchor is located in the peptide colored in red (a.a. 278-289). H2B-ubiquitin is shown in pale green and DNA is shown in pale blue.

To further validate the interactions between DOT1L and nucleosome, we performed hydrogen/deuterium exchange mass spectrometry (HDX-MS). We incubated DOT1L alone, DOT1L_Nuc and DOT1L_Nuc_H2B-Ub in a D2O buffer for 10, 100, 1000, and 10000 seconds. The degrees of hydrogen/deuterium exchanges were analyzed by mass spectrometry (Supplemental Fig. 5). Although we were not able to analyze the HDX-MS for histones due to the limited data quality and low peptide coverage, HDX-MS experiments also revealed that the deuterium uptakes were significantly decreased in several peptides in DOT1L upon nucleosome binding. These peptides include the peptide (_291_RVVELSPLKG_301_) near the hydrophobic patch interacting with the ubiquitin, and the peptide (_278_RINSRNLSDIGT_289_) including the arginine anchor (Figs. 3D and 3E). The deuterium uptake of the arginine anchor peptide is significantly decreased in DOT1L_Nuc compared with DOT1L alone, and further decreased in DOT1L_Nuc_H2B-Ub. These data suggest that the H2B-ubiquitination reorient DOT1L to have tighter interaction with the H2A/H2B acidic patch via the arginine anchor. In addition, the deuterium uptake of the hydrophobic peptide (a.a. 291-301) was further decreased upon H2B-ubiquitination.

These HDX-MS data also support our cryo-EM structure as well as the mutagenesis studies showing that DOT1L recognizes H2B-Ub using the hydrophobic patch including the C-terminal helix, and the H2A/H2B acidic patch using the arginine anchor.

### DOT1L binding destabilizes the nucleosome structure

Unexpectedly, the cryo-EM structures show that DNA is detached from the histone octamer and that the degree of the detachment is greater in DOT1L_Nuc_H2B-Ub compared to DOT1L_Nuc (Fig. 4A). Furthermore, the first helix of histone H3 and the C-terminus of histone H2B were concomitantly disordered with DNA being detached (Fig. 4B). These results suggest that the DOT1L binding to the nucleosome might destabilize the nucleosome structure. Consistent with these observations, a recent study showed that DOT1L bound nucleosome is more readily remodeled by CHD1 ATP-dependent chromatin remodeler than nucleosome alone (Lee et al. 2018).

**Figure 4.**
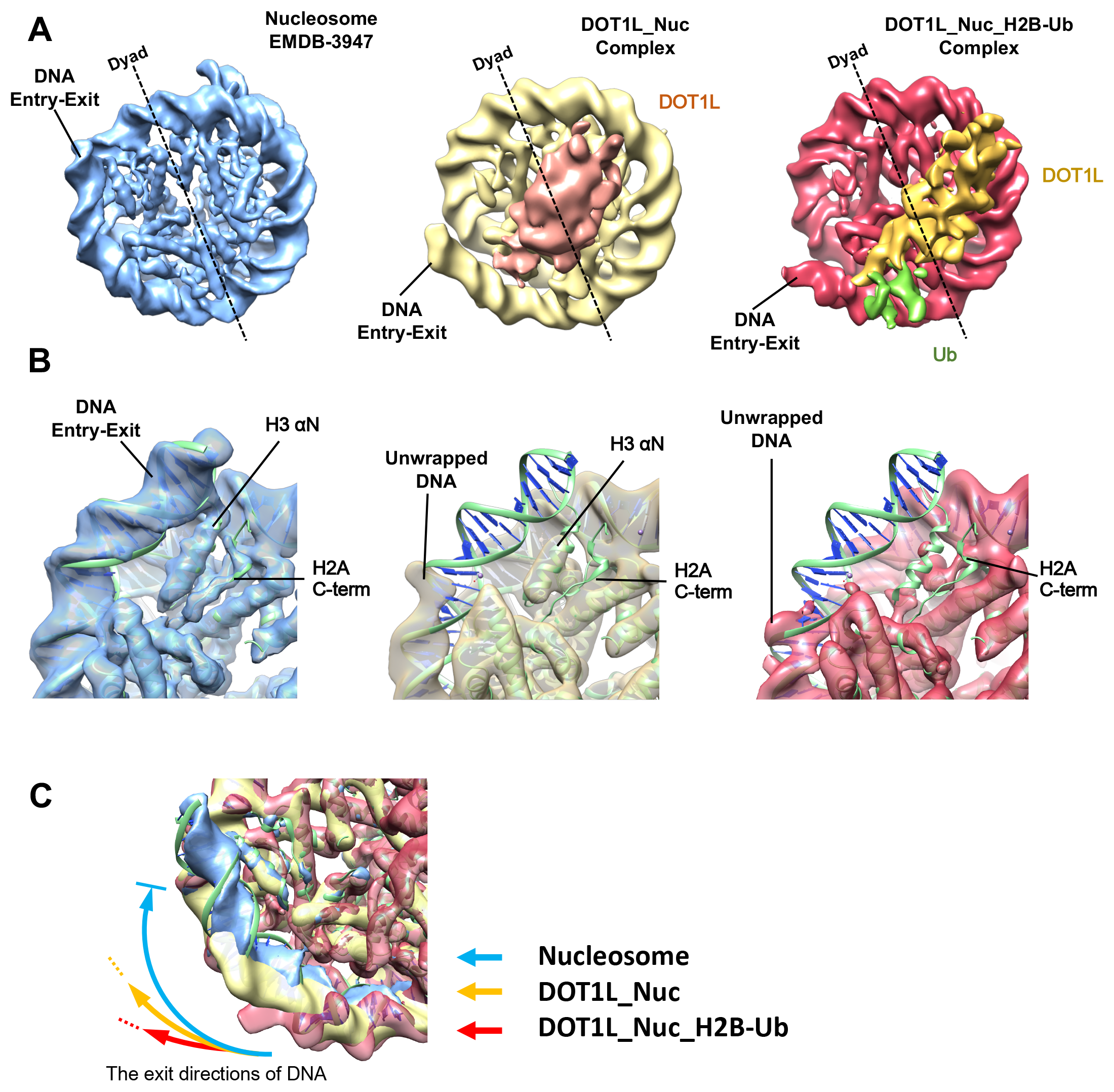
DOT1L mediated nucleosome destabilization. (A) DOT1L binding to nucleosome leads to detachment of DNA and disorder of the first helix of histone H3 and the C-terminus of histone H2B. (A) Cryo-EM maps comparing the position of the DNA exit and entry sites in the canonical nucleosome (EMDB-3947), DOT1L_Nuc and DOT1L_Nuc_H2B-Ub. The DNA at the exit/entry are progressively detached from DOT1L_Nuc to DOT1L_Nuc_H2B-Ub. (B). Detailed view near the DNA exit/entry sites and the histone H3 and histone H2A, which are disordered in DOT1L complexes. (C) Comparison on the exit directions of DNA in the canonical nucleosome, DOT1L_Nuc and DOT1L_Nuc_H2B-ub.

To examine whether the binding of DOT1L destabilizes the nucleosome structure, we analyzed the detachment of DNA from histones using single-molecule FRET experiments. We labeled Cy3 at H2A_T120C_ and Cy5 at the 5’ end of DNA proximal to Cy3 as a FRET pair (Supplemental Fig. 6). The unmodified Nuc or Nuc_H2B-Ub containing the FRET pair was immobilized with biotins conjugated at the other end of DNA on a streptavidin coated surface (Fig. 5A). DOT1L with SAM was then injected and the FRET changes were monitored (Fig. 5B). Upon DOT1L injection, the Cy5 signal slightly increased indicating DOT1L binding to nucleosomes. After initial DOT1L binding, FRET down-spikes were clearly observed, which indicates the detachment of DNA from histones.

**Figure 5.**
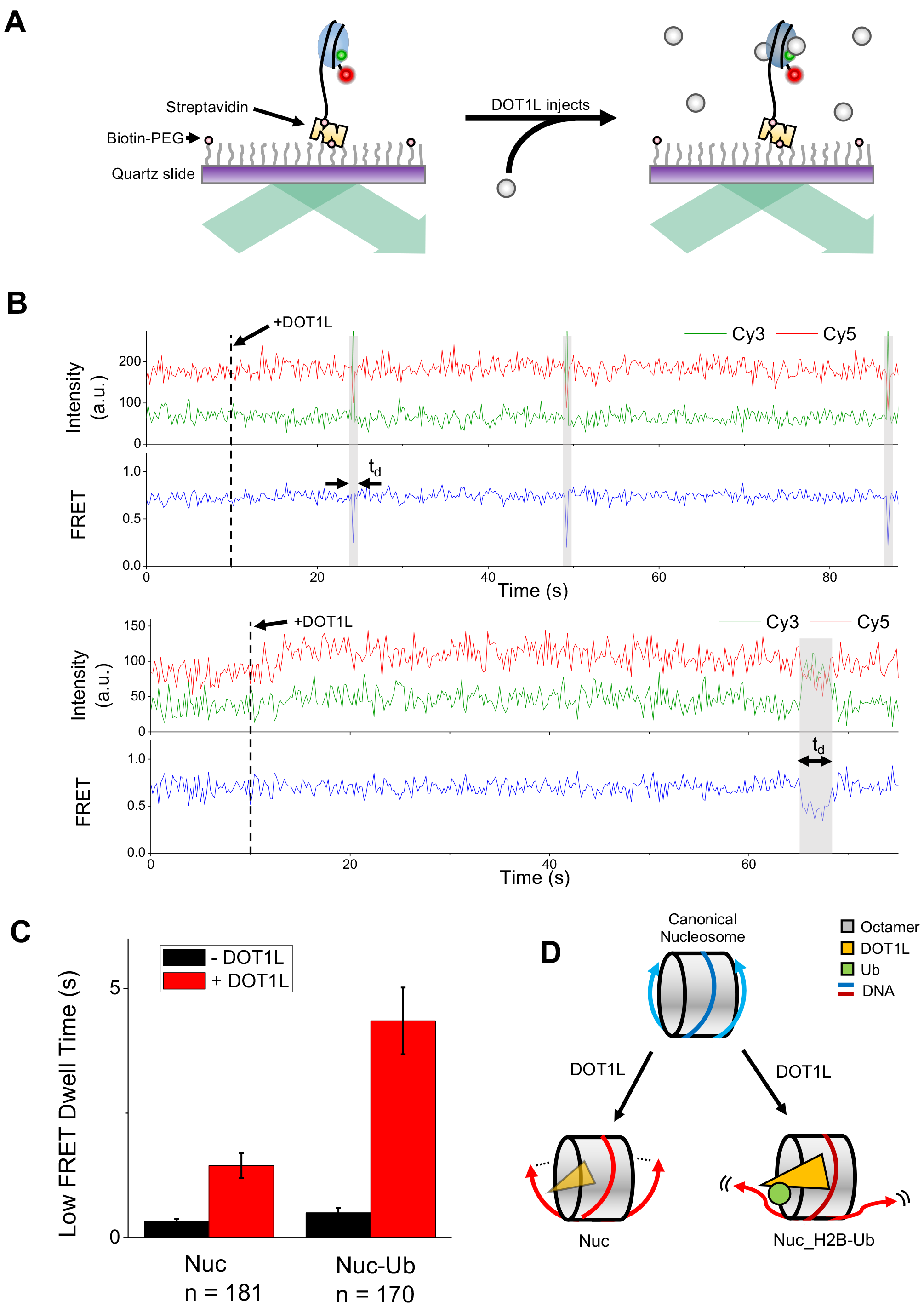
single-molecule FRET showing the destabilization of nucleosome upon DOT1L binding. (A) Experimental scheme. Nucleosomes labeled with a FRET-pair (green circle for Cy3 and red circle for Cy5) were immobilized on a microscope slide. DOT1L was added while single-molecule fluorescence signals were observed. (B) Representative fluorescence intensity (green for Cy3 and red for Cy5) and corresponding FRET (blue) time traces of unmodified nucleosome (top) and ubiquitinated nucleosome (bottom). DOT1L was injected at 10 seconds (dashed lines and black arrows). The increase of Cy5 signal after Dot1L addition indicates the binding of Dot1L to a nucleosome. DNA detachment (FRET down spike) with duration td (gray region) was more clearly observed after Dot1L binding. (C) Low FRET dwell times of unmodified and ubiquitinated nucleosomes in the absence (black) and presence (red) of Dot1L. The data are shown as means ± SEM of five independent experiments. The number of molecules used for the analysis were 181 and 170 for Nuc and Nuc-Ub, respectively. (D) A schematic model for DOT1L mediated nucleosome destabilization.

We then measured and compared the dwell times (t_d_) of the low FRETs, which indicates the duration period that the DNA stays is being detached from histones (Fig. 5C). In Nuc_H2B-Ub (Nuc-Ub), DOT1L binding increased the dwell time (t_d_) of the low FRET state approximately 9-fold compared with nucleosome alone, while in unmodified Nuc (Nuc) DOT1L binding slightly increased the dwell time (t_d_) of the low FRET state. These data clearly show that the DNA detachment lasts substantially longer in Nuc_H2B-Ub than unmodified Nuc upon DOT1L binding. These results from the single-molecule FRET analysis are consistent with our cryo-EM structures showing that the DNA is further detached from histones in DOT1L_Nuc_H2B-Ub than DOT1L_Nuc.

Overall, our single-molecule FRET combined with the cryo-EM structures suggest that DOT1L binding to nucleosomes together with the ubiquitination of H2B greatly destabilize the nucleosome structure. These data demonstrate the molecular basis of the non-catalytic function of DOT1L, which destabilizes nucleosome structure (Figs. 4C and 5D). Furthermore, these results add another layer of complexity to the cross-talk where H2B ubiquitination regulates the DOT1L mediated nucleosome destabilization in addition to histone H3 Lys79 methylation. At this moment, it is not clear how DOT1L binding induces the detachment of DNA with the concomitant disorder of the parts of histones. There are no substantial conformational changes in histones between Nuc alone and Nuc bound to DOT1L although we cannot rule out that the conformational change is too small to be observed at the current resolution. Nonetheless, our single-molecule studies combined with the structures implicate that DOT1L has nucleosome destabilizing acitivty as well as histone methyltransferase activity, and that both activities are affected by H2B ubiquitination.

The results presented here provide the first structural insights into the recognition of H2B ubiquitinated nucleosome by DOT1L. DOT1L recognizes H2B-ubiquited nucleosome via the hydrophobic C-terminal helix for the ubiquitin and the basic anchors for the acidic patches. Furthermore, this work provides the structural evidence of DOT1L-mediated nucleosome destabilization that is enhanced by H2B-ubiquitination. These data implicate the complexity of a cross-talk between H2B-ubiquitinylation and DOT1L activity.

## Materials and Method

### Reconstitution of nucleosomes

N-terminal His-tag ubiquitin_G76c_ (His-Ub) was cloned into a modified pET28a and expressed in BL21(DE3) pLysS *E.coli* cells. The cells expressing the His-ub were lyzed and the cell debris were cleared by centrifugation at 18,000 rpm for 1 hr. His-ub was captured with Ni-NTA resins (Qiagen) and eluted with a buffer containing 500mM NaCl, 50mM Tris-HCl pH 8.0, 10mM imidazole, 10mM β-mercaptoethanol (bME), and 0.1mM imidazole. The eluted His-Ub was dialyzed against a 50mM ammonium acetate pH 4.5 buffer for overnight and purified with HiTrap SP ion exchange chromatography (GE Healthcare). The fractions containing His-Ub were dialyzed into 1 mM acetic acid and the protein was lyophilized for later use. All the *Xenopus laevis* histone H2A, H2B, H2B_k120c_, H3, and H4 were expressed in BL21 (DE3) pLysS E.coli cells and purified according to the published protocol (Luger et al. 1999). Chemically crosslinking between His-Ub_G76c_ and H2B_k120c_ were performed as described in Morgan et al. (Morgan et al. 2016). Reconstitution of nucleosomes was performed with the histones and 601 DNA via the salt dialysis method (Luger et al. 1999).

### DOT1L purification

Wild-type DOT1L catalytic domain (1-416 amino acid) was expressed as a GST-fusion forms. The protein was expressed in BL21 (DE3) RILP *E.coli*. The cell expressing GST-DOT1L were lyzed and the cell debris was cleared with centrifugation at 18,000 rpm for 1 hr. To remove non-specifically bound nucleic acid from the protein, final 0.2% polyethylene imine (PEI) precipitation was performed in the presence of 1 M NaCl to dissociate nucleic acid from the protein. After PEI precipitation, the proteins were collected with 65% ammonium precipitation. The proteins were resuspended and applied to Glutathione Sepharose 4B resins (GE Healthcare). The proteins was eluted and the GST tag was cleaved with HRV3C protease. DOT1L without GST was further purified with a HiTrap SP ion exchange column followed by a Superdex 200 (GE Healthcare) size exclusion chromatography column equilibrated with a buffer containing 300mM NaCl, 50mM Tris-HCl pH8.0, and 2mM DTT. The mutants were purified similarly to the wild-type except that GST was not removed for the HMTase assay.

### Cryo-specimen preparation

To obtain stable complexes of DOT1L and nucleosomes, we adopt Gradient Fixation (GraFix) (Stark 2010). For DOT1L and nucleosome complexes, nucleosome and DOT1L were mixed in 1:5 ratio in 30 mM NaCl, 20 mM HEPES-HCl pH 7.5, 1 mM EDTA, 1mM DTT, 0.5 mM S-Adenosyl Methionine (SAM). The complexes were separated by ultra-centrifugation at 270,000 rpm, 18 hr using a Beckman SW55 rotor in a condition of 5-20% sucrose gradient in the presence of 0-0.2% Glutaraldehyde. 3μl aliquot from the fractions containing the complexes were applied to glow-discharged QuantiFoil™ 2/2, blotted for 3-6 sec, and then plunge-frozen in liquid ethane using a Vitrobot (FEI).

### Cryo-EM image processing

All data were collected at the SciLifeLab, Stockholm using a Titan Krios 300keV with a K2 Summit direct detector. For the DOT1L and unmodified nucleosome complex, a total 2,923 micrographs were collected as 20-frame movie with a pixel size of 1.06 Å/pixel and 4.7 e^-^/Å^2^/sec dose rate. All images were motion-corrected with MotionCor2 (Zheng et al. 2017) and CTFs were calculated by Gctf (Zhang 2016) implemented in Relion2.1. (Kimanius et al. 2016). From 1,301 selected good micrographs, 368,289 particles were picked and subjected to several rounds of 2D-and 3D classifications. The initial model was generated using EMAN2 (Tang et al. 2007). Finally, 21,299 particles were used for final reconstruction. The refined cryo-EM structure showed 7.3 Å resolution as determined by the gold standard FSC at 0.143 criterion. The detailed scheme for the processing is shown in Supplementary Fig. 2A. For the DOT1L and H2B-ubiquitinated nucleosome data set, a total 3,831 micrographs were collected as 20-frame movie with a pixel size of 1.06 Å/pixel and 4.7 e^-^/Å^2^/sec dose rate. The motion correction was done with MotionCor2 and CTFs were calculated by Gctf implemented in Relion2.1. A total 430,300 particles from 2,857 selected micrographs were picked and subjected to several rounds of 2D-and 3D classification. The initial model was generated using cisTEM (Grant et al. 2018). Finally, 122,242 particles were used to produce the final cryo-EM structure. According to the gold standard FSC at 0.143 criterion, the final resolution was 6.8 Å. Reconstructions were post-processed in Relion2.1. The local resolutions were analyzed using ResMap (Kucukelbir et al. 2014) (Supplemental Figs. 2C and 3C).

### Modul building

For initial modeling, the crystal structures of the nucleosome core particle, DOT1L, and ubiquitin (PDB 1KX3, 1NW3, and 1UBI, respectively) were docked into the Cryo-EM density using the *Fit in Map* module of UCSF Chimera (Pettersen et al. 2004); each of the crystal structures was treated as a rigid-body during the initial global search with UCSF Chimera. Next, the atomic coordinates of DOT1L and ubiquitin were further optimized by molecular dynamics flexible fitting (MDFF) (Trabuco et al. 2008). We performed two consecutive runs of MDFF for the refinement following the default MDFF protocol, carrying out 200,000 MDFF steps for each run. The flexible fittings converged rapidly in all the MDFF runs. The cryo-EM maps and the pdb coordinates of DOT1L_Nuc and DOT1L_Nuc_H2B-Ub will be deposited to the EMDB and PDB with the accession ID of XXX, XXX, XXX and XXX, respectively.

### Histone methyltransferase assay

2Wild type and mutant DOT1Ls (125 or 250 fmol) were incubated with unmodified or H2B-ubiquitinated nucleosomes (1 pmol) in a condition of 50 mM Tris-HCl pH 9.0, 5 mM MgCl2, 4 mM DTT) in the presence of S-adenosyl methionine SAM[^3^H] (PerkinElmer) at 37°C for 1 hour. The reactions were stopped by SDS-loading buffer and the proteins were separated by 18% SDS-PAGE and transferred to PVDF (Merck) membrane. The membranes were then exposed to an imaging plate for overnight and tritium (^3^H) signals were detected by using FUJI BAS. The band intensities were quantified by using Image J.

### Hydrogen/deuterium exchange mass spectrometry (HDX-MS)

HDX was initiated by mixing 5 μL of 15~20 μM protein samples with 25 μL of D2O buffer (20 mM HEPES, pD 7.5, 30 mM NaCl, 1mM EDTA, and 1mM DTT in D2O) and incubating for 10, 100, 1000, and 10000 s at 4°C. At the indicated time points, each mixture was quenched by adding 30 μL of ice-cold quenching buffer (100 mM NaH2PO4, pH 2.01, 20 mM TCEP, and 0.3 M guanidine hydrochloride). For non-deuterated samples (ND), 5μL of protein samples was mixed with 25 μL of H2O buffer (20 mM HEPES, pH 7.5, 30 mM NaCl, 1mM EDTA and 1mM DTT in H2O) and quenched with 30 μL of ice-cold quench buffer. The quenched samples were digested by passing through an immobilized pepsin column, and the peptide masses were analyzed as described previously (Bang et al. 2018). Peptides from non-deuterated samples were identified with ProteinLynx Global Server (PLGS) 2.4 (Waters, Milford, MA, USA). The following parameters were applied: monoisotopic mass, nonspecific for the enzyme while allowing up to 1 missed cleavage, MS/MS ion searches, automatic fragment mass tolerance, and automatic peptide mass tolerance. Searches were performed with variable methionine oxidation modification. To process HDX-MS data, the amount of deuterium in each peptide was determined by measuring the centroid of the isotopic distribution using DynamX 2.0 (Waters, Milford, MA, USA). The back-exchange level was not corrected because all the data was shown as a comparison of different states. All data were derived from 4 independent experiments, and ANOVA, Bonferroni test and unpaired t-test were performed by SPSS.

### Single-molecule FRET

The nucleosomes used in single molecule FRET experiment were designed to have a 78-bp spacer on the biotinylated side and a 3-bp spacer on the unbiotinylated side (Supplemental Fig. 6). DNA fragments containing 601 nucleosome positioning sequence were generated by PCR as described (Lowary and Widom 1998; Lee et al.2017). Primers with Cy5 at the 5’ end or biotin were purchased from IDT. PCR products were purified by 5% native PAGE gel. Nucleosomes were reconstituted with histone octamer containing Cy3-labeled H2A T120C and Cy5-labeled DNA fragment using salt gradient dialysis. Quartz slides and coverslips were cleaned with piranha solution (mixture of 3:1 concentrated sulfuric acid:30% [v/v] hydrogen peroxide solution), coated with aminopropyl silane first, and then with mixture of PEG (m-PEG-5000, Laysan Bio) and biotin-PEG (biotin-PEG-5000, Laysan Bio). A flow cell was assembled by combining a coverslip and a quartz slide with double-sided adhesive tape(3M). For convenient buffer exchange, polyethylene tubes (PE50; Becton Dickinson) were connected to the flow cell. Nucleosome substrates were immobilized on a PEGylated surface using biotin-streptavidin conjugation. Single-molecule FRET experiments were performed in an imaging buffer (10mM Tris-HCl pH 8.0, 100 mM KCl, 3 mM MgCl2, 5 mM NaCl) containing a gloxy oxygen scavenging system (a mixture of glucose oxidase (1 mg/ml, Sigma), catalase (0.04 mg/ml, Sigma), glucose (0.4% (w/v), Sigma), and Trolox (2 mM, Sigma)). Single-molecule fluorescence images were acquired at a frame rate of 10 Hz using a home-built prism-type total internal reflection fluorescence microscope equipped with an electron-multiplying charge coupled device camera (Ixon DV897; Andor Technology). Cy3 and Cy5 were alternately excited with 532nm and 633nm lasers using the ALEX(Alternative Laser Excitation) technique (Kapanidis et al. 2005). Experimental temperature was maintained at 30^Ộ^C using a temperature control system (Live Cell Instruments). Data acquisition and FRET trace extraction were done using homemade programs written in LabView (National Instruments) and IDL (ITT), respectively. The intensity traces of Cy3 and Cy5 were analyzed using a custom made Matlab(MathWorks) script. The FRET efficiencies were calculated from the donor (ID) and acceptor (IA) fluorescence intensities as EFRET = IA/(IA + ID).

## Acknowledgements

We thank the Song lab members for helpful discussion. We thank Drs. Carol Soomin Cho and Soung-Hun Roh for critical comments on the manuscript. We also thank the staffs at the SciLife Lab, Stockholm for the data collection. This work is supported by grants (NRF-2016R1A2B3006293, NRF-2016K1A1A2912057) to J.S., a grant (NRF-2012R1A5A2A28671860) to K.Y.C. from National Research Foundation (NRF) of Korea, and a grant (2016-03810) from Swedish Research Council to H.H. The computing resource was supported by Global Science experimental Data hub Center (GSDC), Korea Institute of Science and Technology Information (KISTI) (RF-2018R1A6A7052113). S.J. is a recipient of the Global Fellowship (2016H1A2A1908806) by Korea National Research Foundation.

## Author contributions

S.J and J.S. conceived the idea, and S.J., T.Y., H.H. and J.S. determined the cryo-EM structures. C.K. and S.H. performed the single-molecule FRET experiments, H.Y. and K.Y.C. performed the HDX-MS experiments, and S.J.K. performed the molecular dynamics flexible fitting. S.J. and J.S. wrote the manuscript and all authors reviewed the data and the manuscript.

