## Supplemental Information for "Structural basis of recognition and destabilization of histone H2B ubiquitinated nucleosome by DOT1L histone H3 Lys79 methyltransferase"

Supplemental Table 1. Cryo-EM data collection and validation statistics.

| | DOT1L_Nuc | DOT1L_Nuc_H2B-Ub | Comment |
|---|---|---|---|
| **Sample Preparation** | | | |
| Grid | Quantifoil 2/2 | Quantifoil 2/2 | Quantifoil |
| Cryo-specimen freezing | Vitrobot 4 | Vitrobot 4 | FEI |
| **Data Collection** | | | |
| Electron microscope | FEI Titan KRIOS G3i | FEI Titan KRIOS G3i | 300 KeV |
| Detecting device | Gatan K2 summit | Gatan K2 summit | Electron counting |
| Total Electron Exposure ($e^-/Å^2$) | 37.28 | 33.6 | |
| Dose rate ($e^-/Å^2$/sec) | 4.7 | 4.8 | |
| Number of frames in each movie | 20 | 20 | |
| Exposure time (sec) | 8 | 7 | |
| Defocus range (μm) | -1.8 to -4 | -1.8 to -4 | |
| Pixel size (Å) | 1.06 | 1.06 | |
| **Image processing** | | | |
| Processing software | EMAN2.0/RELION2.1 | CisTEM/RELION2.1 | |
| Initial particle images (no.) | 368,289 | 430,300 | |
| Final particle images (no.) | 21,229 | 122,242 | |
| Symmetry imposed | C1 | C1 | |
| Map resolution (Å) | 7.3 | 6.8 | Gold Standard FSC (0.143) |
| Map-sharpening *B*-factor ($Å^2$) | -100 | -300 | |
| **Model Validation** | | | |
| Bonds (RMSD) | | | |
| Length (Å) | 0.028 (506) | 0.031 (626) | |
| Angles ( °) | 2.034 (416) | 2.164 (508) | |
| Ramachandran plot (%) | | | |
| (Favored/Allowed/Outlier) | 96.10/2.60/1.30 | 96.00/2.70/1.30 | |
| CC between map and model | 0.65 | 0.73 | |

**Supplemental Fig. 1. Ubiquitinated H2B production and protein purification**

(A) SDS-PAGE followed by Coomassie staining of unmodified H2B and ubiquitinated H2B. (B) SDS-PAGE followed by Coomassie staining of unmodified nucleosome and H2B-ubiquitinated nucleosome (C) SDS-PAGE followed by Coomassie staining of wild-type and mutant DOT1Ls.

**Supplemental Fig. 2. Structural determination of DOT1L unmodified nucleosome complex**

(A) A detailed scheme for image process of DOT1L_Nuc. (B) A local resolution on the map by ResMap (top) and Euler angle distribution of the particles used for the reconstruction (bottom). (C) FSC curve for DOT1L_Nuc indicates 7.3 Å resolution by the gold standard FSC at 0.143 criterion.

**Supplemental Fig. 3. Structural determination of DOT1L H2B-ubiquitinated nucleosome complex**

(A) A detailed scheme for image process of DOT1L_Nuc_H2B-Ub. (B) A local resolution on the map by ResMap (top) and Euler angle distribution of the particles used for the reconstruction (bottom). (C) FSC curves for DOT1L_Nuc_H2B-Ub indicates that 6.8 Å resolution by the gold standard FSC at 0.143 criterion.

**Supplemental Fig. 4. Cryo-EM maps of DOT1L_Nucleosome complexes**

(A) Cryo-EM maps of DOT1L and unmodified nucleosome complexes. (B) Cryo-EM maps of DOT1L and H2B-ubiquitinated nucleosome complex

**Supplemental Fig. 5.  Peptide coverage for DOT1L in HDX-MS**

The dark-blue boxes indicate the peptide identified from HDX-MS.

**Supplemental Fig. 6. Singe-particle FRET**

(A) DNA sequences used for single-molecule FRET experiments. The DNA fragments consist of 601 nucleosome positioning sequence (black) with the 3bp linker on the exit side and the 78bp on the entry side (blue). The Cy5 labeling position is marked in red. (B) FRET histogram and labeling scheme of the nucleosome. The histogram shows

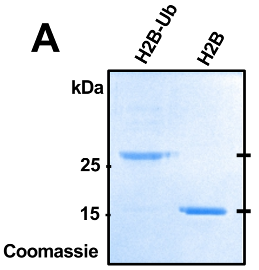
A
H2B-Ub
H2B
kDa
25
15
Coomassie

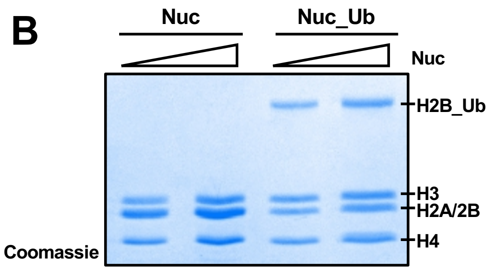
B
Nuc
Nuc_Ub
Nuc
H2B_Ub
H3
H2A/2B
H4
Coomassie

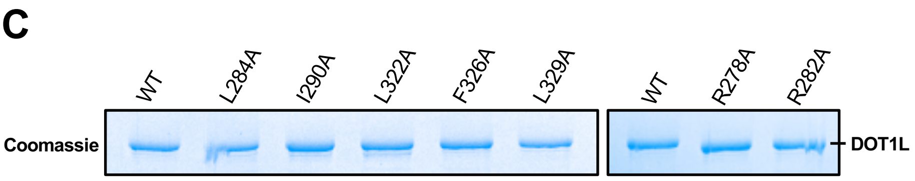
C
WT
L284A
I290A
L322A
F326A
L329A
WT
R278A
R282A
Coomassie
DOT1L

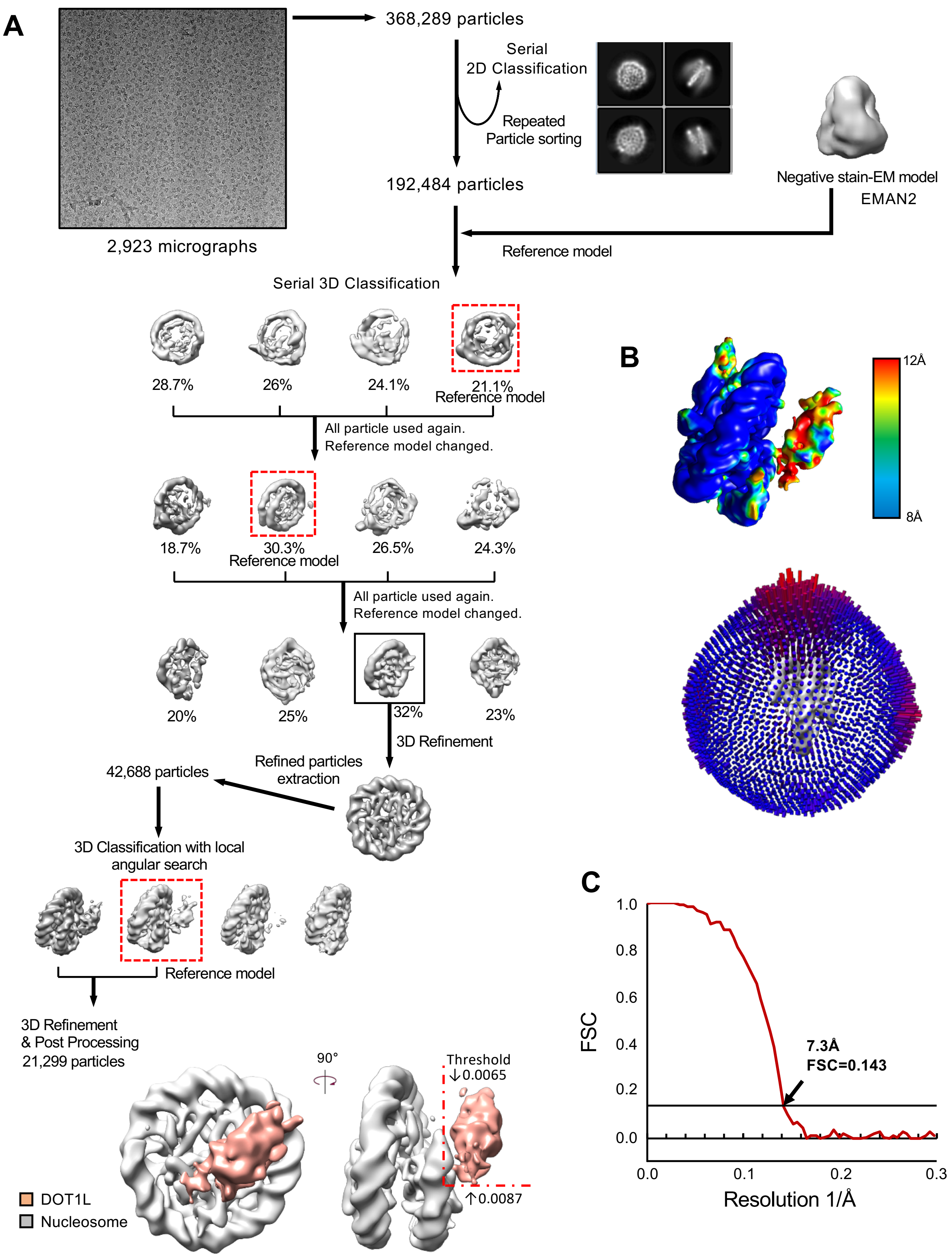
A
368,289 particles
Serial
2D Classification
Repeated
Particle sorting
192,484 particles
2,923 micrographs
Negative stain-EM model
EMAN2
Reference model
Serial 3D Classification
28.7%
26%
24.1%
21.1%
Reference model
All particle used again.
Reference model changed.
18.7%
30.3%
Reference model
26.5%
24.3%
All particle used again.
Reference model changed.
20%
25%
32%
23%
3D Refinement
Refined particles
extraction
42,688 particles
3D Classification with local
angular search
Reference model
3D Refinement
& Post Processing
21,299 particles
90°
Threshold
↓0.0065
↑0.0087
DOT1L
Nucleosome
B
12Å
8Å
C
FSC
1.0
0.8
0.6
0.4
0.2
0.0
7.3Å
FSC=0.143
0.0
0.1
0.2
0.3
Resolution 1/Å

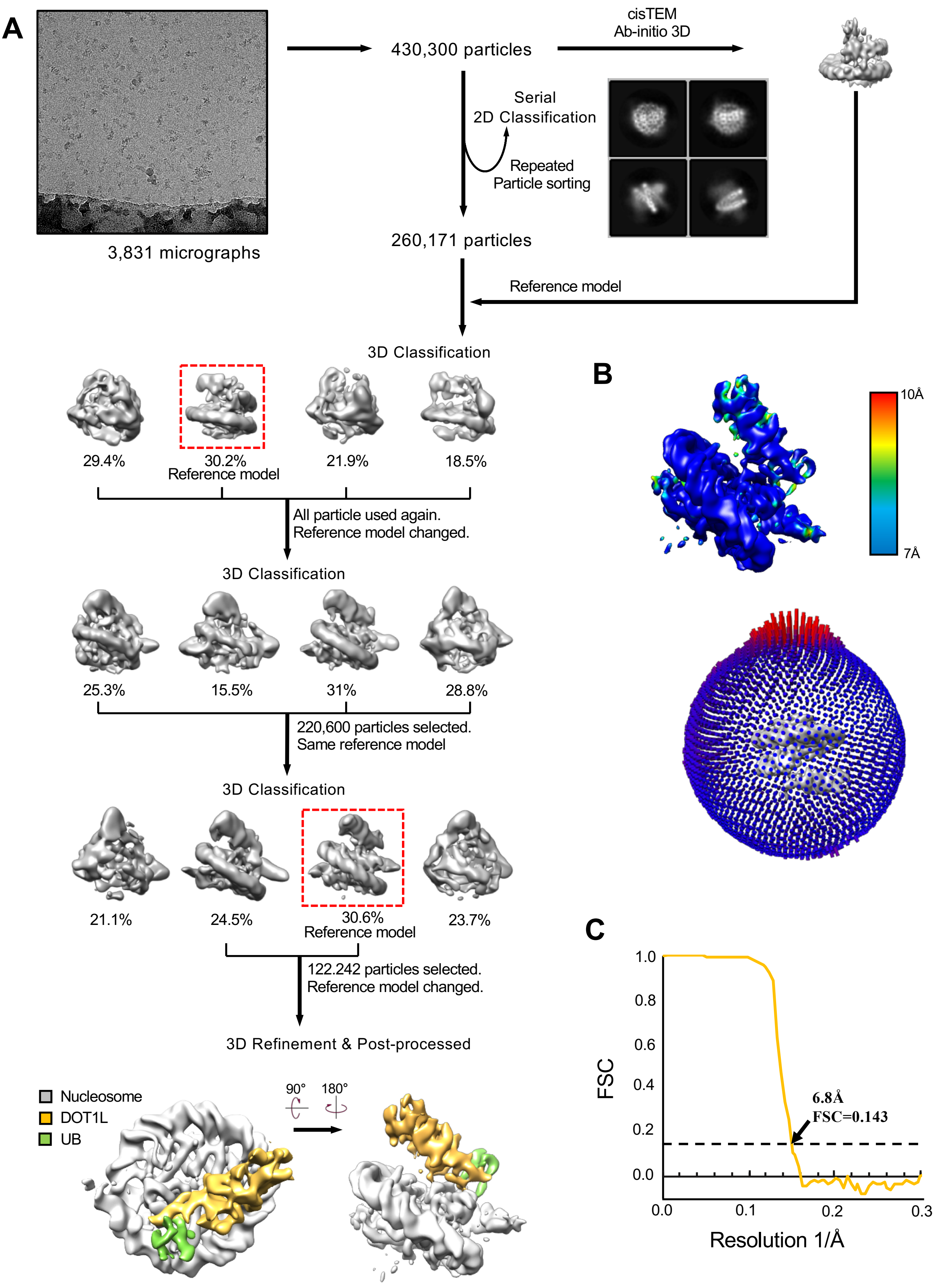
A
3,831 micrographs
430,300 particles
cisTEM
Ab-initio 3D
Serial
2D Classification
Repeated
Particle sorting
260,171 particles
Reference model
3D Classification
29.4%
30.2%
Reference model
21.9%
18.5%
All particle used again.
Reference model changed.
3D Classification
25.3%
15.5%
31%
28.8%
220,600 particles selected.
Same reference model
3D Classification
21.1%
24.5%
30.6%
Reference model
23.7%
122.242 particles selected.
Reference model changed.
3D Refinement & Post-processed
Nucleosome
DOT1L
UB
90°
180°
B
10Å
7Å
C
FSC
6.8Å
FSC=0.143
1.0
0.8
0.6
0.4
0.2
0.0
0.0
0.1
0.2
0.3
Resolution 1/Å

**A**

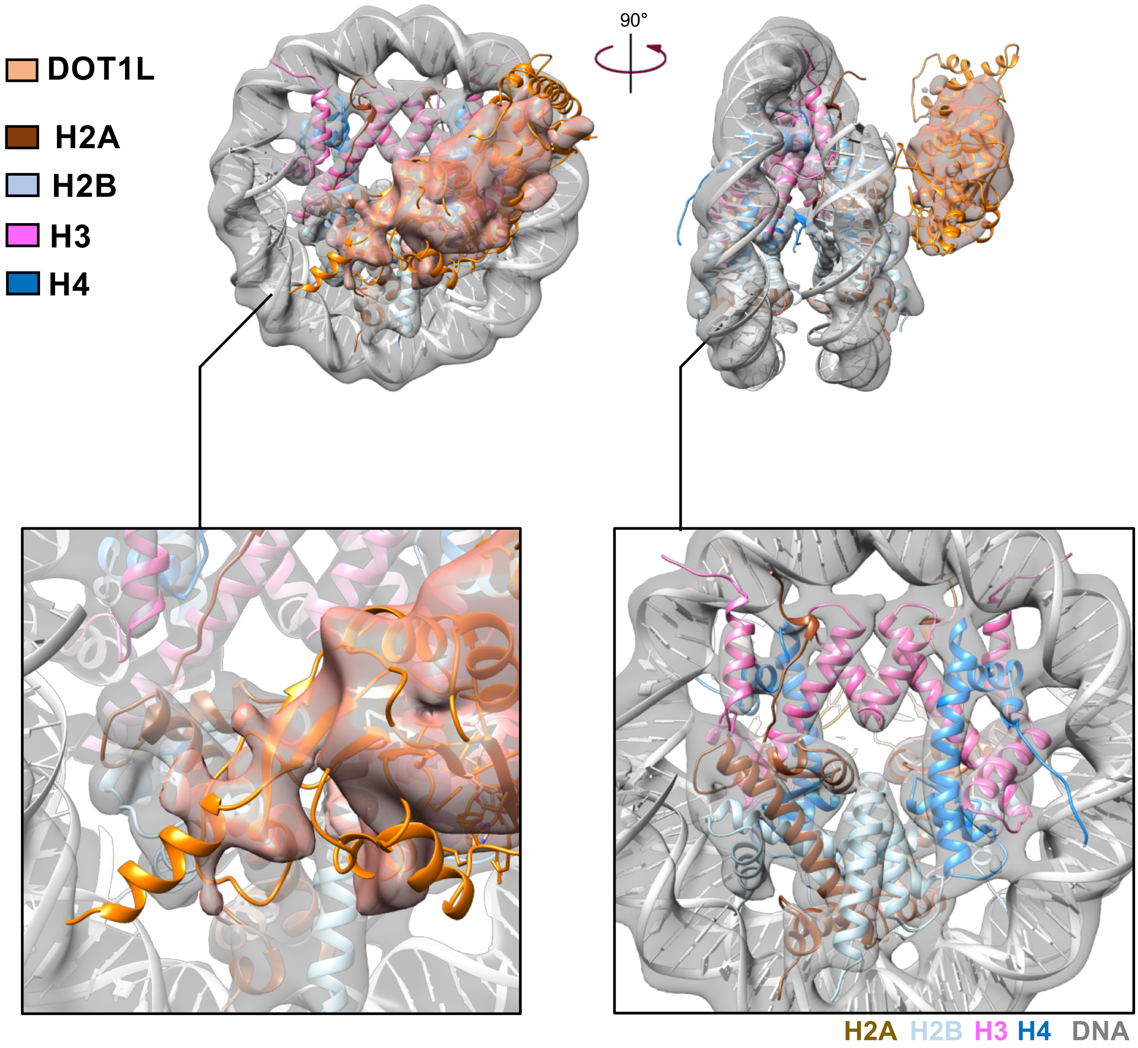

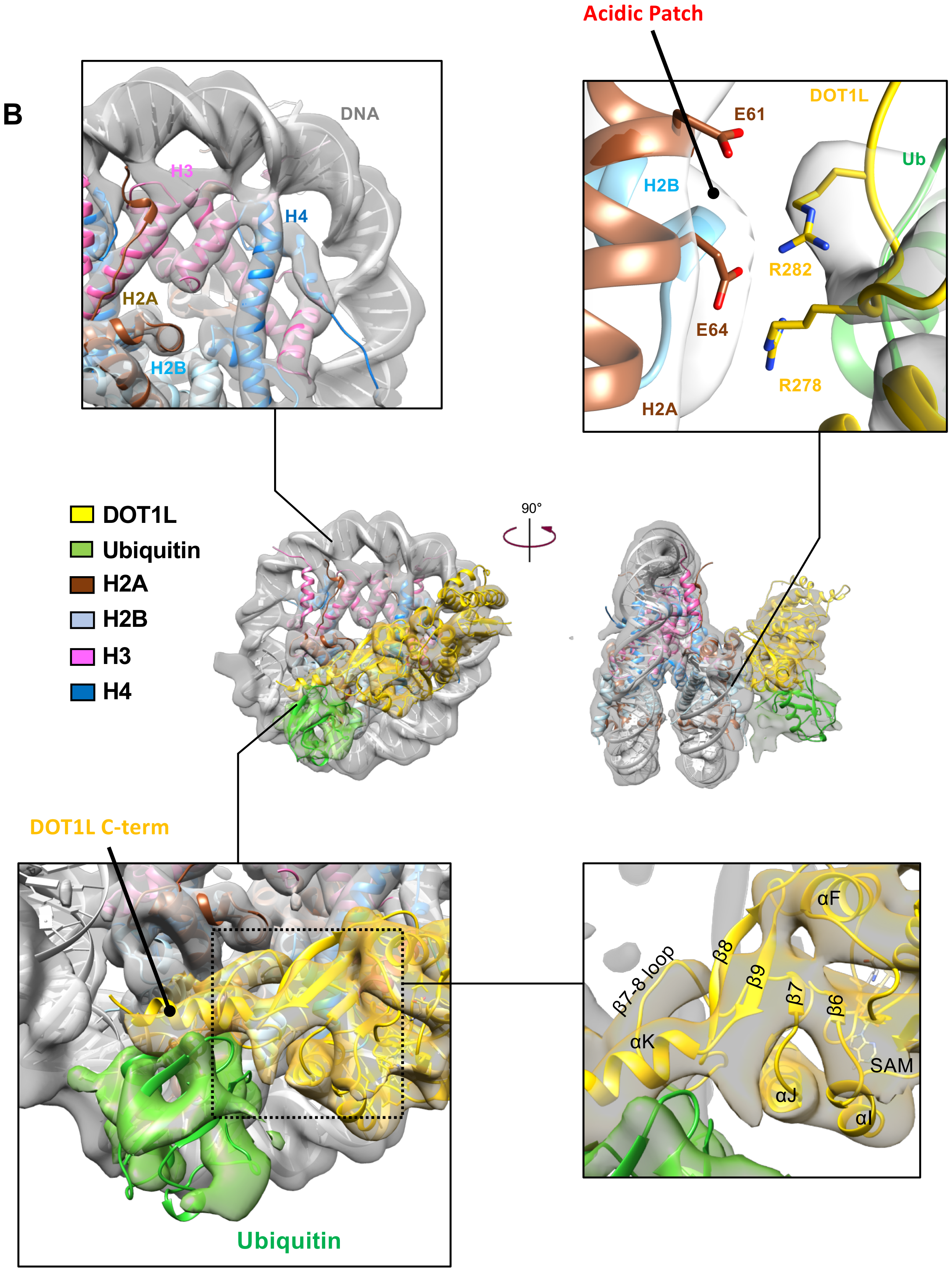
B
Acidic Patch
DNA
H3
H4
H2A
H2B
E61
DOT1L
Ub
H2B
R282
E64
R278
H2A
DOT1L
Ubiquitin
H2A
H2B
H3
H4
90°
DOT1L C-term
Ubiquitin
αF
β8
β7-8 loop
β9
β7
β6
αK
SAM
αJ
αI

**Jang et al., Supplemental Fig. 5**

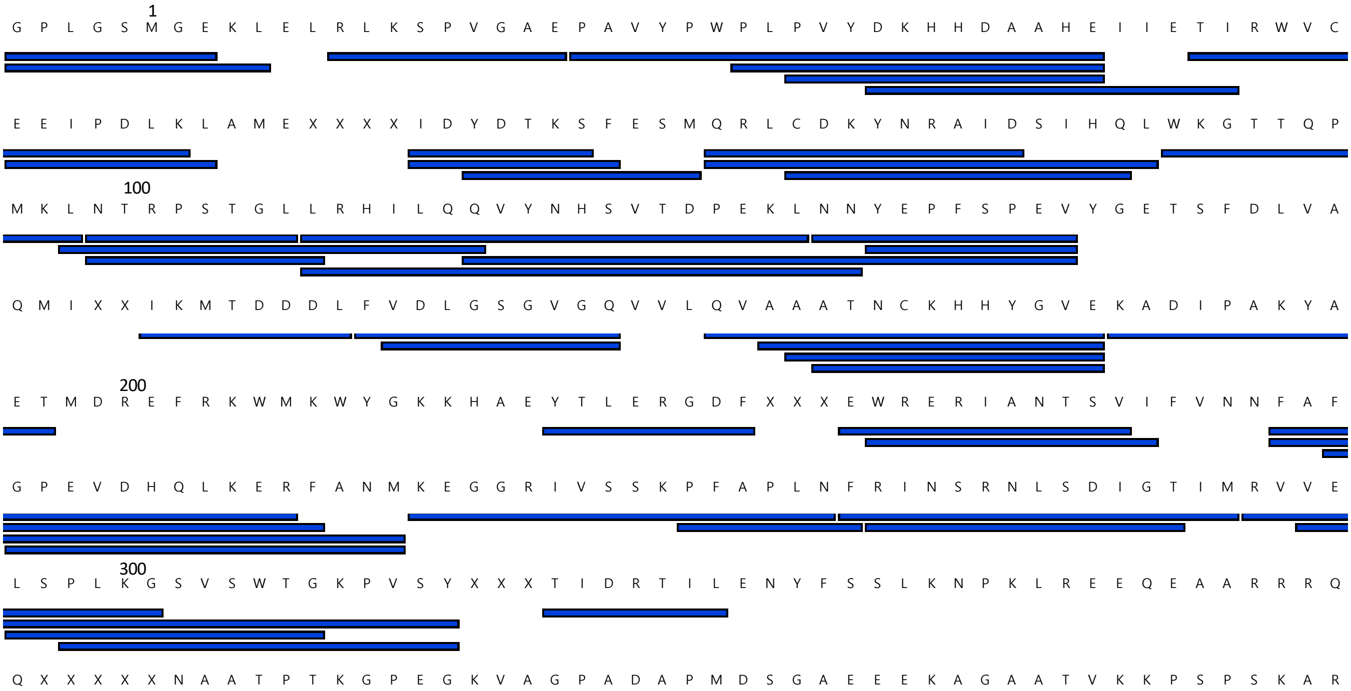


(A) HDX-MS of DOT1L. The dark-blue boxes indicate the peptide covered from HDX-MS.

## A

GCCCT GGAGA ATCCC GGTCT GCAGG CCGCT CAATT GGTCG TAGAC AGCTC
TAGCA CCGCT TAAAC GCACG TACGC GCTGT CCCCC GCGTT TTAAC CGCCA
AGGGG ATTAC TCCCT AGTCT CCAGG CACGT GTCAG ATATA TACAT CCTGT
GCATG TATTG AACAG CGACC TTGCC GGTGC CAGTC GGATA GTGTT CCGAG
CTCCC ACTCT AGAGG ATCCC CGGGT ACC

(A) DNA sequences used for single-molecule FRET experiments. The DNA fragments consist of 601 nucleosome positioning sequence (black) with the 3bp linker on the exit side and the 78bp on the entry side (blue). The Cy5 labeling position is marked in red.

## B

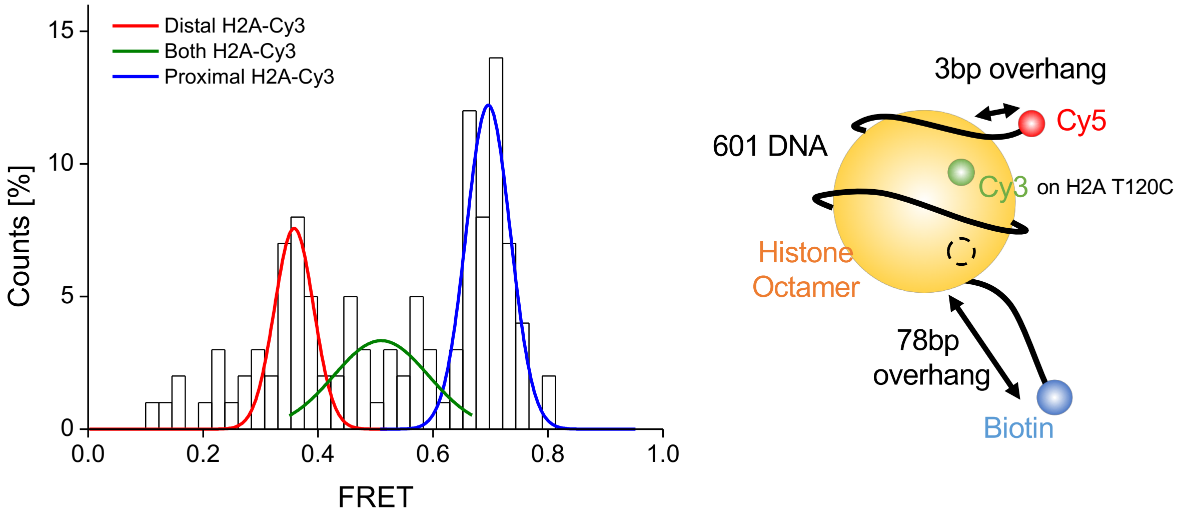


(B) FRET histogram and labeling scheme of the nucleosome. The histogram shows three FRET peaks. The low and high FRET correspond to species labeled at the distal and proximal H2A, respectively. The middle FRET corresponds to doubly-labeled species. We selected only high FRET species for data analysis.

three FRET peaks. The low and high FRET correspond to species labeled at the distal and proximal H2A, respectively. The middle FRET corresponds to doubly-labeled species. We selected only high FRET species for data analysis.s
